# Evolution of drought resistance strategies following the introduction of white clover (*Trifolium repens* L.)

**DOI:** 10.1101/2024.09.03.611023

**Authors:** Brandon T. Hendrickson, Caitlyn Stamps, Courtney M. Patterson, Hunter Strickland, Michael Foster, Lucas J. Albano, Audrey Y. Kim, Paul Y. Kim, Nicholas J. Kooyers

## Abstract

**Background and Aims:** Success during colonization likely depends on growing quickly and tolerating novel and stressful environmental conditions. However, rapid growth, stress avoidance, and stress tolerance are generally considered divergent physiological strategies.

**Methods:** We evaluate how white clover (*Trifolium repens*) has evolved to a divergent water regime following introduction to North America. We conduct RNAseq within a dry-down experiment utilizing accessions from low and high latitude populations from native and introduced ranges, and assess variation in dehydration avoidance (avoidance of wilting) and dehydration tolerance (ability to survive wilting).

**Key Results:** Introduced populations are better at avoiding dehydration, but poorer at tolerating dehydration than native populations. There is a strong negative correlation between avoidance and tolerance traits and expression of most drought-associated genes exhibits similar tradeoffs. Candidate genes with expression strongly associated with dehydration avoidance are linked to stress signaling, closing stomata and producing osmoprotectants. However, genes with expression linked to dehydration tolerance are associated with avoiding excessive ROS production and toxic bioproducts of stress responses. Several candidate genes show differential expression patterns between native and introduced ranges, and could underlie differences in drought resistance syndromes between ranges.

**Conclusions:** These results suggest there has been strong selection following introduction for dehydration avoidance at the cost of surviving dehydration.

## INTRODUCTION

Species introductions are occurring at a drastically increasing rate due to globalization and human commerce (Seebens *et al*. 2017; Bonnamour *et al*. 2021). Determining how and why some introduced species become invasive is a persistent goal for biologists that has had relatively limited success (Kolar and Lodge 2001; Pyšek and Richardson 2007). Introduced species almost certainly encounter some level of novel abiotic or biotic environments during initial colonization, and must both establish and spread to become invasive. While historically neglected as an important evolutionary force during invasion, selection is commonly observed during successful invasions - especially for human commensal species that are introduced repeatedly and/or have high genetic variation among colonists (Huey 2000; Maron *et al*. 2004; Colautti and Lau 2015; Battlay *et al*. 2024).

Selection is key for many of the hypotheses that describe why some species become successful invaders. Stress resistance and phenotypic plasticity have frequently been hypothesized as key attributes for successful invasive species (Richards *et al*. 2006; Gewing *et al*. 2019). Individuals that are able to avoid or tolerate stress can better establish and spread in novel environments.

Key to this hypothesis is that selection acts to remove less stress resistant individuals from the populations. Such selection could also occur only in certain areas of the introduced range where selection pressures are strongest. Alternatively, hypotheses like the Evolution of Increased Competitive Ability (EICA; Blossey & Notzold, 1995) suggest that introduced plants are removed from key selective pressures in the native range following colonization and the strongest selection pressure is simply for rapid growth rate and high fecundity to outcompete native species.

One of the most important selection pressures for plants is water availability and associated drought events (Siepielski *et al*. 2017). Three different non-mutually exclusive strategies are hypothesized to evolve in environments with low or inconsistent water availability: drought escape, dehydration avoidance and dehydration tolerance (Ludlow 1989; Kooyers 2015; Volaire 2018). Plants that have evolved to escape drought grow rapidly, reproduce quickly, and finish their life cycles prior to or immediately following drought stress. Plants that are dehydration avoiders actively evade drought conditions with physiological or morphological adaptation or acclimation responses to limit desiccation and thus wilt at a lower soil volumetric water content. While dehydration avoidance can be a constitutive strategy where certain individuals may have high succulence, more trichomes, higher water-use efficiency, or lower stomatal density, it also manifests as a response to reduced water availability by lowering stomatal conductance, raising water-use efficiency, and producing osmoprotectants for osmotic adjustment. Plants with dehydration tolerant strategies limit water-loss during drought conditions by limiting vegetative growth, allocating resources to root growth, and producing antioxidants to limit cellular damage from desiccation. These responses allow plants to recover following intense drought. A common experimental design to tease apart variation in drought strategies between populations and/or species is conducting dry-down experiments (e.g. Des Marais *et al*., 2014; Bouzid *et al*., 2019; FitzPatrick *et al*., 2023). In these experiments, variation in time to first flower can be considered a drought escape metric, differences in soil relative water content at wilting can be considered a measure of dehydration avoidance, and the ability to recover following wilting is a measure of dehydration tolerance (Bouzid *et al*. 2019).

While drought strategies are not mutually exclusive, negative correlations between some strategies are expected due to physiological constraints. For instance, the rapid growth expected for drought escape requires opening stomata in order to increase photosynthetic rate (Geber and Dawson 1990; Dudley 1996). However, open stomata increase transpiration and lower water use efficiency – effectively neutralizing any dehydration avoidance strategy. Yet, the evidence for this tradeoff is equivocal and seemly context dependent (McKay *et al*. 2003; Kooyers *et al*. 2015). Correlations between dehydration avoidance and tolerance strategies are less explored, and are often viewed as compatible responses (Volaire 2018). However, resource allocation tradeoffs could be hypothesized as the functions necessary for maintaining water balance in a limited water environment are not the same resources necessary for surviving and recovering from dehydration.

Although the molecular signals underlying the perception and integration of water stress are becoming better resolved, variation in drought-related responses is rarely understood at the molecular level (VanWallendael *et al*. 2019). For instance, it is largely unknown whether variation in drought resistance arises through differences in sensitivity or perception of drought, integration of signals, or re-wiring of connections within gene-regulatory networks. Molecular responses to low water availability center around the activation of the abscisic acid (ABA) dependent and ABA-independent signal transduction pathways in response to disruption in water potential relative to the surrounding soil and/or atmosphere (Raghavendra *et al*. 2010; Dar *et al*. 2017; Aslam *et al*. 2022). Increases in ABA drive a signal cascade leading to guard cell regulation and stomatal closure (Hsu *et al*. 2021). Notably, production of ABA also induces gene regulatory cascades that lead to increases in antioxidants and osmoprotectants such as proline, trehalose, and raffinose family oligosaccharides (Strizhov *et al*. 1997; Lunn *et al*. 2014). The interactions of ABA with other plant hormones (specifically antagonistic interactions with gibberellic acid and ethylene), lead to cessation of shoot growth and/or increased root growth (Wilkinson and Davies 2010). Not surprisingly, this complex web of gene regulatory cascades has substantial overlap – for instance, products of the SNF1-related kinases 2 (*SnRK2*) gene family integrate growth rate changes, promote stomatal closure, and increase osmoprotectants (Yoshida *et al*. 2015; Belda-Palazón *et al*. 2020). A key goal for the plant science community is determining how these networks are manipulated to create the diversity of responses observed within and among species (Kesari *et al*. 2012; Des Marais *et al*. 2014). Invasive species with divergent responses between introduced and native areas can provide key natural experiments for examining the evolution of drought resistance.

White clover (*Trifolium repens*) is perennial plant that is native to Eurasia but has been distributed across the world as a cover crop and forage legume during the past 300 years (Kjærgaard 2003). Despite relatively recent introductions, there is little decline in genetic diversity in the introduced ranges of white clover (Wu *et al*. 2021) and extensive evidence for selection and adaptation following introduction (Kooyers and Olsen 2012, 2013; Wright *et al*. 2018, 2021; Santangelo *et al*. 2022; Battlay *et al*. 2024). While white clover is most often associated with temperate habitats, there is extensive variation in aridity across both the native and introduced ranges, and water availability seems to be a key selective pressure (Kooyers *et al*. 2014). Escape, avoidance, and tolerance responses have been noted by different groups. Drought escape via more annualized life history is favored in lower latitude populations in North America (Wright *et al*. 2021). Dehydration avoidance may be favored in these same populations as clines in cyanogenic glucosides, potentially key sources of nitrogen during drought conditions, have evolved following introduction to several different continents. In each introduced region, the proportion of plants possessing cyanogenic glucosides increases in more arid areas (Kooyers & Olsen 2013, 2014). Finally, agricultural breeders frequently target persistence following drought as a key trait for selection in agricultural lines (Marshall *et al*. 2001; Jahufer *et al*. 2002).

However, differences in drought resistance strategies have not been systematically examined across native and introduced populations.

Here we use a manipulative dry-down experiment to examine how populations across the native and invasive ranges of white clover resist drought. Specifically, we examine the relative degree of dehydration avoidance and tolerance among populations from low and high latitude populations in the native and invasive range using volumetric water content at wilting as a measure of dehydration avoidance and ability to recover following wilting as a measure of dehydration tolerance. We also examine differential expression between well-watered and dry down plants at a timepoint when the dry down lines begin to experience drought stress. We use this data to address four questions: First, do populations from introduced areas or from more arid areas have more substantial drought resistance strategies? The stress tolerance hypothesis suggests that introduced plants should be better stress tolerators, but EICA may suggest the opposite. Second, are there tradeoffs that exist among dehydration avoidance and tolerance strategies? That is, are plants that are good dehydration avoiders also poor dehydration tolerators? Third, do native and introduced populations exhibit different patterns of gene expression in response to early drought stress? That is, are differentially expressed genes between well-watered and drought treatments also variable among ranges and populations? Fourth, how do expression differences control variation in dehydration avoidance and tolerance strategies? Together these experiments provide insight on the molecular mechanisms underlying the evolution of drought resistance across a rapid invasion.

## MATERIALS AND METHODS

We performed a manipulative experiment to examine variation among drought strategies and the transcriptomic drivers of such variation within and among the native and introduced ranges of white clover, *Trifolium repens*. We planted seed collected from 3-4 populations from low latitude and high latitude in the native and invasive range (14 total populations; Fig. S1; Innes *et al*., 2022). Seeds for each population had been pooled from >25 different maternal lines. For each population, we used latitude and longitude to extract WorldClim2 variables (Fick and Hijmans 2017) and annual aridity and evapotranspiration data from CGIAR (Zomer *et al*. 2008).

Seeds were scarified and planted into 4” pots filled with PRO-MIX LP-15 soil pre-saturated with water. Pots were placed in AR-66L2 growth chambers (Percival Scientific; Perry, IA, USA) and misted daily to promote germination. Growth chambers were set to a constant 22°C with a 14hr:10hr day:night cycle. Germinants were randomized within well-watered and dry-down treatments (well-watered = 26 individuals, dry-down = 25 individuals; Table S1). Plants were rotated within chambers every three days to minimize microenvironmental variation. Plants were grown for six weeks to accumulate above and below ground biomass. At six weeks, all pots were saturated with water by bottom-watering. Number of leaves on each plant were counted and one pot was excluded from further analysis because it was substantially larger than the other plants. Plants in the well-watered (control) treatment received periodic watering to saturation. Plants in the dry-down treatment did not receive additional water. Each day, we assessed volumetric water content in each pot using a SMT150T soil moisture meter (Dynamax; Houston TX, USA). Well-watered and dry-down treatments functioned as expected with separation in volumetric water content (VWC) between treatments after control plants were resaturated on day 9 (Fig. S2)

To examine dehydration avoidance, we recorded the day and the soil moisture at which each plant in the dry-down treatment wilted (hereafter, “wilt VWC”). To determine dehydration tolerance, we determined the propensity to recover following wilting (hereafter “wilt survival”). Forty-eight hours after wilting, we resaturated the pots of each wilted plant by bottom watering for an extended period of time (∼1hr). We then resurveyed plants for survival three days (72hrs) after resaturation. Survival was measured as a discrete variable (yes/no) with any revived green tissue or new growth counting as survival.

### Manipulative Experiment Statistics

All statistics were conducted in R v4.2.2 (R Foundation for Statistical Computing, Vienna, Austria) with univariate linear models (LM) and generalized linear models (GLM) implemented with the lm() or glm() functions, respectively. First, to examine whether plant size influenced either dehydration avoidance or tolerance, we examined associations between either wilt VWC and wilt survival against leaf number at the beginning of the experiment. Neither wilt VWC or survival was significantly associated with leaf number (Fig. S3). Models below produce qualitatively similar results whether or not leaf number at day 1 is included as a covariate.

We examined whether variation in dehydration avoidance and tolerance was associated with range (native/introduced), latitude (low/high) or an interaction using generalized linear models. Specifically, we modeled wilt VWC and wilt survival as response variables within a GLM with range, latitude, and range:latitude interaction as factors. Wilt VWC was modeled with a gaussian distribution with an identity link and wilt survival was modeled with a binomial distribution with a logit link. Statistical significance was determined through ANOVA using a type II sum of squares implemented with the *Anova()* function (*car* library; Fox *et al*., 2013) Latitude across continental ranges may not be strongly related to water availability or aridity of populations and thus may not be the main factor driving variation in drought strategies. To examine whether variation in drought strategies was related to differences in precipitation or aridity, we examined associations between wilt VWC and wilt survival with annual aridity index, potential evapotranspiration, and coefficient of variation in annual precipitation (bio15).

Distributions and links of models as well as assessment of statistical significance were the same as the above models. Leaf number at day 1 of the experiment was included as covariate in each analysis as there were weak associations between each climatic variable and initial plant size (Fig. S4).

To explore potential tradeoffs between dehydration avoidance and tolerance, we examined associations between wilt VWC and wilt survival using a GLM (binomial distribution, logit link). Ability to recover from wilting was the response variable and volumetric water content at wilting the predictor variable. Significance was tested with an ANOVA as above and model fit was assessed using the *adjR2()* function (glmToolbox package; Vanegas *et al*., 2024) .

### RNAseq Analysis

We used an RNAseq analysis to examine differential expression underlying variation in drought strategies. During the manipulative experiment, leaf tissue from two healthy adult leaves was flash frozen in liquid nitrogen ten days after the dry down treatment began from plants in both the well-watered and dry-down treatment. Tissue sampling was done during a two-hour period in the afternoon with order of collection randomized within each treatment. Total RNA was extracted with Direct-zol RNA Miniprep (Zymo Research; Irvine, CA, USA). RNAseq libraries were constructed with QuantSeq 3’ mRNA-Seq FWD prep kit (Lexogen; Vienna, Austria) using an input of ∼415ng total RNA/sample following manufacturers protocols. Two samples had low RNA concentrations and we followed Lexogen’s modified procedure for low input samples.

Four samples from the manipulative experiment did not have RNAseq libraries constructed due to loss of labels during shipping or near-zero read counts (RNAseq N = 47; Table S2). All individuals were barcoded and multiplexed in a single tube. We sequenced this library on a single HiSeqX lane (PE 150bp reads) through Novogene (Sacramento, CA, USA). A second round of multiplexing and sequencing was conducted for a subset of samples that had low data yield the first round, and files from both rounds were subsequently merged. Raw sequenced reads per individual averaged 37,241,613 reads (SD 16,534,847; Table S2) Fastp v.023.4; (Chen *et al*. 2018) was used to trim adapters and poly-a tails. Microbial RNA contamination was removed by aligning each sample to all fungal and bacterial assembly genomes within the NCBI database using bowtie2 v2.5.1(Langmead and Salzberg 2012). The bacterial and fungal genome database was indexed with the 2.2.4 release of bowtie2. The remaining reads that did not align to the contamination database were presumed to be white clover RNA reads. A transcriptome was created from the *T. repens* reference genome (Santangelo *et al*. 2023) using gffread v0.12.7 (Pertea and Pertea 2020). A full decoy-aware transcriptome was constructed to mitigate specious mapping of reads that occur from unannotated genomic loci that are similar to the annotated transcriptome, and a mapping-based index was then constructed using a k-mer hash over k-mers of length 31 using Salmon v1.10.2 (Patro *et al*. 2017). Read mapping to the transcriptome and quantification was then performed with Salmon v1.10.2 in mapping-based mode using the *T. repens* transcriptome and RNA sequences. There was a clear issue with DNA contamination as only 3.83% of reads per individual aligned to the transcriptome (SD 1.0%; Table S2); however, this should not bias our comparative analysis (differential expression or transcript abundance-phenotype associations) that rely on read counts of each transcript.

### Quantifying expression across treatment, range, and latitude

We used DESeq2 (Love *et al*. 2014) to test for differences in transcript abundance between dry-down and well-watered treatment groups, between the North American and European range, and between high and low latitude populations. We controlled for the VWC at the time of tissue collection by treating it as a covariate in the DESeq2 model (i.e., Transcript ∼ Treatment*Latitude*Range + SoilVWC). False discovery rate (FDR) for each gene was calculated and a transcript was then categorized as differentially expressed if FDR was < 0.1.

Concordance of gene expression responses to drought between ranges and latitudes was determined by comparing the coefficient estimates of protected one-way ANOVAs. That is, we conducted ANOVAs for each gene found to be differentially expressed between the well-watered and dry-down treatments with relative expression as the response variable and either range or latitude as the predictor variable.

All transcripts were used for weighted correlation network analysis (WGCNA) to determine gene modules and co-expression networks within the WGCNA v1.72-5 library (Langfelder and Horvath 2008). To achieve homoskedasticity amongst transcript abundance, the variance stabilization transformation from the fitted dispersion-mean relations was calculated with DESeq2, which was then used to transform transcript count data by dividing by the size factor.

Normalized transcript counts were then analyzed with WGCNA using a soft-thresholding power of 9 and assuming that biological networks follow a scale-free structure (Barabási and Albert 1999). The sets of genes within each resulting module were then cross referenced with the ‘biological process’ gene ontology terms (GO; Ashburner *et al*., 2000) of *Arabidopsis thaliana* and used for gene enrichment analysis with an FDR cutoff of 0.1. Fold enrichment, –log10(FDR), and number of genes for each GO term was then calculated. Genes belonging to the same module were then compared to the Kyoto Encyclopedia of Genes and Genomes (KEGG; Kanehisa & Goto, 2000) to visualize expression pathways.

### Identifying genes associated with wilt VWC and wilt survival

GLMs were constructed to quantify the correlation of transcript abundance with either wilt VWC or wilt survival. First, transcripts of drought treatment samples were standardized to acquire relative expression for each gene. Univariate GLMs included relative expression as the response variable and either wilt VWC (gaussian distribution, identify link) or wilt survival (binomial distribution, logit link) as predictor variables with VWC at the start of the experiment as a covariate. Significance was then tested using ANOVA as above. To determine the set of phenotypically significant genes that are co-expressed, genes significantly associated with each phenotype were compared to the previously calculated WGCNA network. We also determined the location of these genes within the white clover genome (Santangelo *et al*. 2023) via *blastn* (Altschul *et al*. 1990).

Finally, we then determined which of the genes associated with phenotype also were differentially expressed between ranges or across latitudes to identify genes likely to be involved in adaptation following introduction or to divergent climatic conditions. We conducted GLMs modeling relative expression as a function of traits and either range or latitude. Specifically relative expression was the response variable while trait value (wilt VWC or wilt survival), spatial group (range or latitude), and the interaction between trait:group were included as predictors. Genes with relative expression that is explained by trait:group interaction are good candidates for producing the phenotypic variation observed in nature.

## RESULTS

There was substantial variation in dehydration avoidance and tolerance throughout the native and introduced ranges of white clover. Wilt VWC was lower in North American populations than in European populations (F1,23 = 4.8, p = 0.03; Fig. 1A) indicating greater drought avoidance in introduced populations. However, the effect size of this difference was low as the difference in mean wilt VWC was only 1.3% lower water content at wilting in the North American populations. There were no differences among low and high latitude populations or any interaction between introduction and latitude impacting dehydration avoidance (Table S3).

**Fig. 1.**
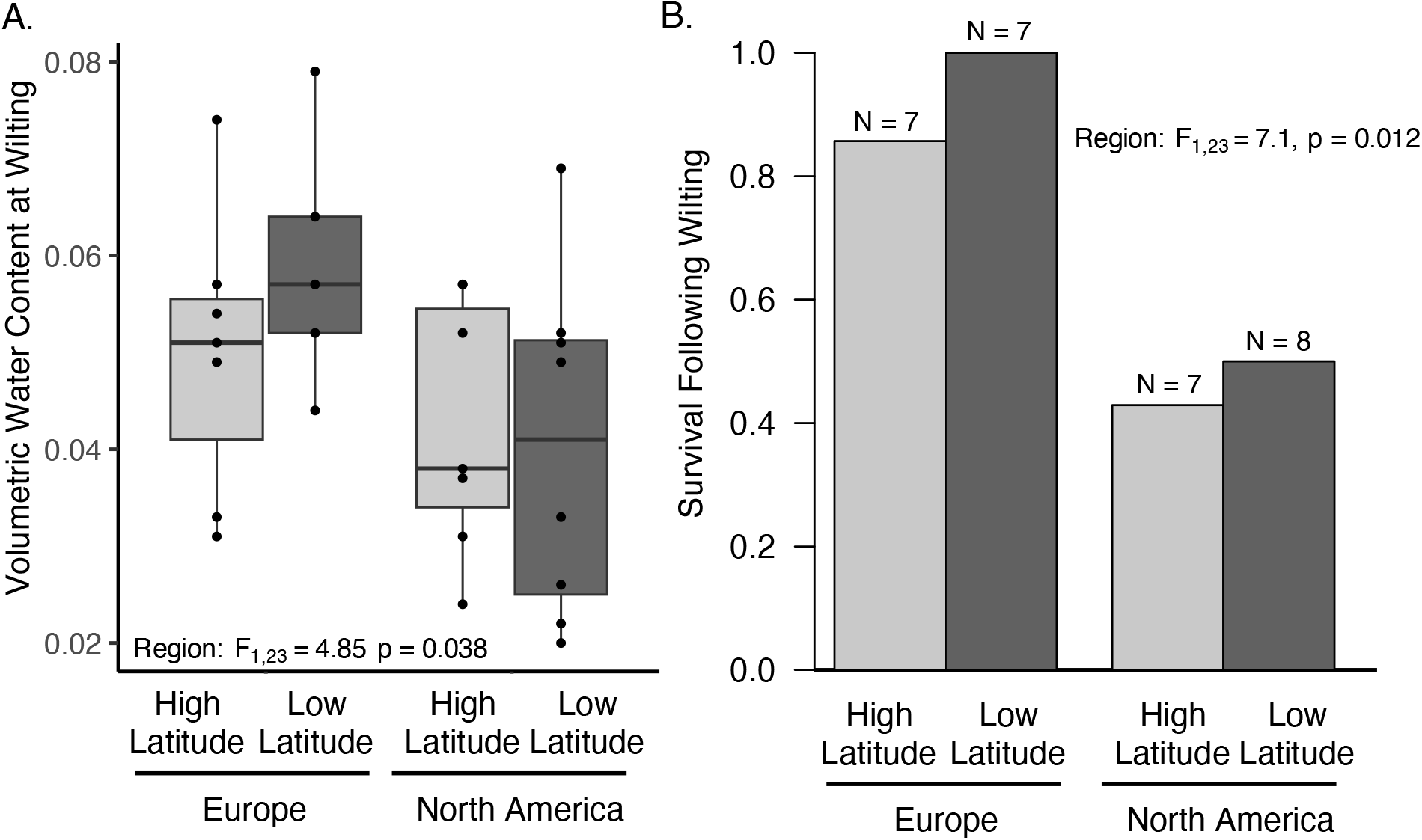
Variation in drought strategies between native and introduced populations of white clover. (A). Boxplots of variation in wilt VWC within low and high latitude population in the native (European) and introduced (North American) regions. Each point is an individual. Box edges represent the interquartile range, the center line in the box is the median, and the whiskers represent 1.5 times less or greater than the interquartile range. (B). Barplots identify the proportion of individuals that survived the wilting treatment from each latitude:region contrast. Numbers above the bars are the number of individuals surveyed.

Dehydration tolerance also differed among native and introduced ranges with native regions able to better survive wilting (Ξ^2^ = 7.09, p = 0.012; Fig. 1B). All but one of the plants from the native range were able to recover following wilting (11/12 plants), but only 46.7% (7/15) plants recovered from the introduced North American range. There were no differences among low and high latitude populations or any interaction between introduction and latitude impacting dehydration tolerance (Table S2).

There are strong associations between the abiotic environment of site where each line originated and wilt survival, but little correlation between abiotic variables and wilt VWC. Dehydration tolerance was associated with aridity index, with plants from areas with greater annual aridity index being more likely to survive wilting (Ξ^2^ = 6.2, p = 0.013; Fig. S5). This pattern was driven by the low latitude populations from the native range that all recovered following wilting. These low latitude populations, largely from Spain, had relatively high evapotranspiration, low annual precipitation, and higher variation in precipitation. Dehydration avoidance was not significantly associated with any abiotic variables (Fig S5.; Table S4).

There was a strong tradeoff between dehydration avoidance and tolerance. Plants that wilted at lower VWC were less likely to recover following wilting (Adj. r^2^ = 0.53, p = 0.005; Fig. 2). The four plants wilting at the lowest volumetric water contents all did not survive following wilting. Moreover, six of the seven plants with the lowest wilt VWC also did not survive wilting.

**Fig. 2.**
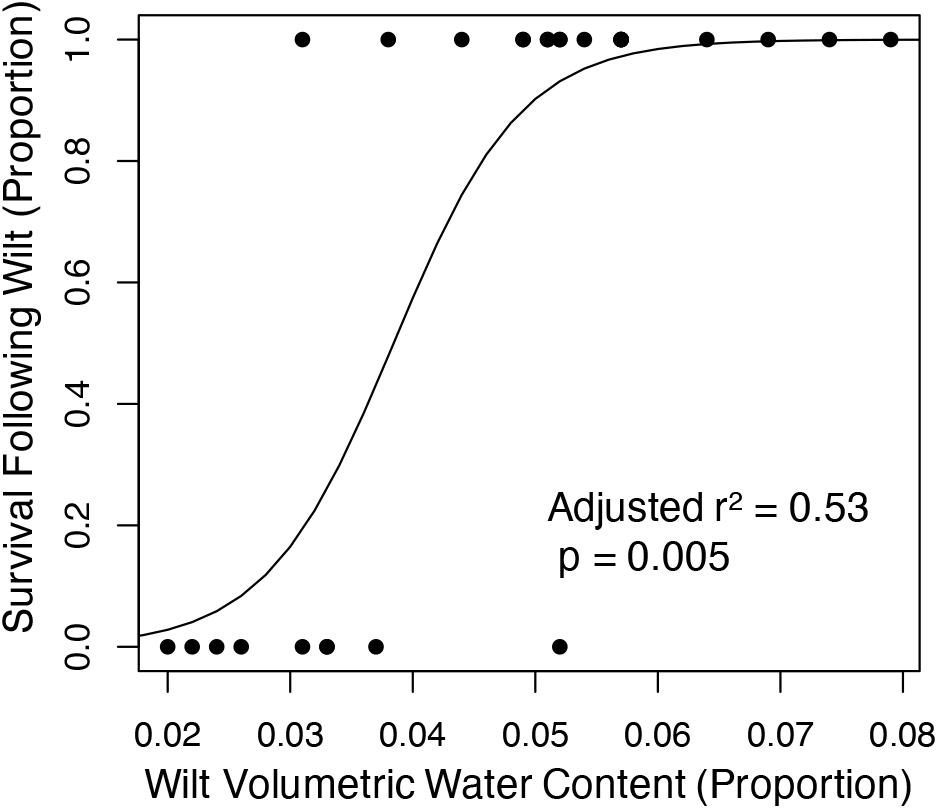
Phenotypic tradeoff between dehydration avoidance and tolerance. Lower wilt VWC indicates greater dehydration avoidance while higher wilt survival indicating greater dehydration tolerance.

### Differential expression across the native and introduced ranges

Many genes were differentially expressed between dry down and well-watered treatments, high and low latitude populations, and native and introduced ranges (Table S5). Across predictor variables, there were 19,344 differentially expressed transcripts out of 86,723 sequenced transcripts corresponding to 2,253 of 5,001 unique genes. Most differentially expressed transcripts were differentially expressed within all three contrasts (Treatment, Range, Latitude; 35.3%; Fig. S6). An additional 29.4% of transcripts were differentially expressed both in Range and Latitude contrasts, not in the Treatment contrast. A far lower percentage of differentially expressed transcripts were differentially expressed in only one contrast (Latitude: 13.4%; Treatment: 5.6%; Range: 10.0%). The large union between the three contrasts in differentially expressed transcripts indicates different drought responses are likely occurring across latitudinal gradients and ranges.

We examined co-expression between differentially expressed genes to determine how gene expression networks are influenced by drought treatment and vary across range and latitude. Eight modules were discovered from WGCNA including 5.77% of transcripts (Table 1, Table S5). Expression profiles of normalized transcript counts show the “turquoise” module as having elevated transcript counts in samples within the dry-down treatment in contrast to all other modules that exhibit relative uniformity across sample types (Table 1; Fig. S7). Gene enrichment analysis of the “turquoise” module reveals marginal or significant enrichment for many GO terms typically associated with drought resistance including: hyperosmotic stress, proline metabolic process, response to oxygen-containing compound, and response to water deprivation (Table S6). Within the “turquoise” module, 41 of the 53 genes (77.4%) exhibit a significant latitude by range interaction (24 homologues exhibit significantly different range responses between high and low latitudes) – substantially higher than the genes not in modules (18.2%; Table 1). These results suggest that genes within the turquoise module may underlie adaptation to climate across the native and introduce ranges.

**Table 1:** Breakdown of the genes and module associations within phenotypes and across space.

| Module | # Transcripts | Wilt VWC | Wilt Survival | Latitude | Range | Latitude X Range |
| --- | --- | --- | --- | --- | --- | --- |
| No-Module | 45916 (4360) | 2066 (557) | 1201 (336) | 2994 (854) | 1076 (309) | 3895 (875) |
| In-Module | 4337 (662) | 346 (89) | 240 (62) | 603 (169) | 170 (51) | 592 (130) |
| *brown* | 2298 (428) | 158 (50) | 110 (29) | 426 (134) | 97 (28) | 221 (54) |
| *red* | 122 (36) | 8 (1) | 7 (2) | 20 (5) | 5 (1) | 15 (4) |
| *yellow* | 259 (45) | 3 (1) | 7 (1) | 16 (5) | 3 (2) | 50 (9) |
| *grey* | 821 (120) | 52 (6) | 28 (5) | 37 (8) | 34 (11) | 36 (6) |
| *turquoise* | 285 (53) | 86 (28) | 59 (15) | 70 (12) | 8 (0) | 196 (41) |
| *green* | 138 (31) | 5 (1) | 5 (2) | 8 (2) | 8 (4) | 7 (1) |
| *blue* | 324 (86) | 27 (5) | 21 (8) | 19 (8) | 15 (6) | 51 (20) |
| *black* | 91 (20) | 7 (0) | 3 (0) | 7 (1) | 0 (0) | 16 (5) |

Number outside of parentheses refers to the number of transcripts that are associated with the phenotype or differentially expressed across Latitude or Range contrasts. The number within the parentheses refers to gene homologues that are associated with the phenotype or differentially expressed across Latitude or Range contrasts.

### Differentially expressed genes are associated with dehydration avoidance and tolerance

Expression of a large number of genes was associated with wilt VWC and wilt survival (635 and 392, respectively; 198 shared; Table S6). The overlap between this group and genes that are differentially expressed within the dry-down vs. well-watered contrast represent the best candidate genes for impacting dehydration avoidance and tolerance strategies. A promising number of genes are found in this overlapping group – 47 differentially expressed genes were associated with wilt VWC and 17 differentially expressed genes with wilt survival (7 shared between phenotypes). These genes disproportionately were found in the ‘turquoise’ module: 80% of gene homologs for wilt VWC and 50% of gene homologs for wilt survival.

From this initial group, we identified several strong candidate genes by examining genes with extreme effect sizes on each phenotype and are also known drought-associated genes in other species. We first examined genes which were more highly expressed in plants with low wilt VWCs as genes potentially involved in dehydration avoidance. Several genes from the turquoise co-expression network had the greatest negative beta estimates for wilt VWC. These included multiple homologs of *NAC4* and *BCAT2* (Fig. 3AB). Notably, expression of these genes was also strongly associated with death following wilting, i.e. the most negative beta estimates for dehydration tolerance. Several other strong candidate genes had this same strong tradeoff between phenotypes including homologs of *AlaAT2*, *GOLS2*, *SDIR1* and *P5SC1* (Fig. 3CDEF).

**Fig. 3.**
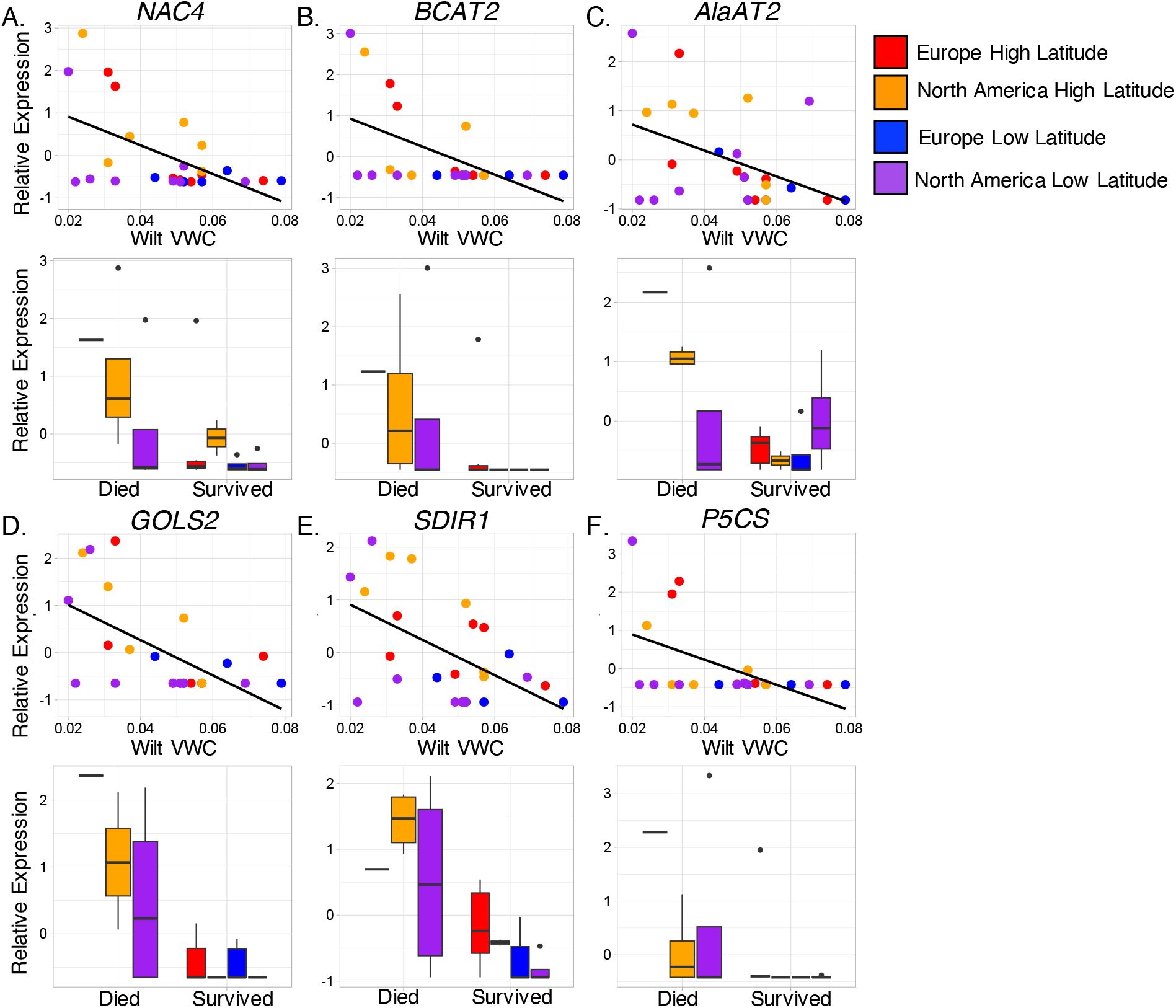
Association between candidate gene expression and dehydration avoidance and tolerance phenotypes. Lower wilt VWC indicates greater dehydration avoidance while higher wilt survival indicating greater dehydration tolerance. Points and bar are colored by latitude and range: low latitude Europe = blue, high latitude Europe = red, low latitude North America = purple, High Latitude North American = orange. Each point represents a different individual. Gene Abbreviations: *NAC4* = nascent polypeptide-associated complex, *BCAT2* = branched chain amino acid transaminase 2, *AlaAT2* = alanine aminotransferase 2, *GOLS2* = galactinol synthase 2, *SDIR1* = Salt and Drought-Induced Ring Finger1, *P5CS* = delta 1-pyrroline-5-carboxylate synthase.

In fact, all transcripts that had significant associations with both wilt VWC or wilt survival had either negative beta for both wilt VWC and wilt survival or positive betas for both traits (α =0.05, 582 transcripts & 170 homologues; Fig. 4). We discuss gene functions in more depth in the discussion, but wilt VWC-associated genes are strongly related to stress signaling via both ABA-dependent and independent pathways, stomatal responses, and producing various osmoprotectants.

**Fig. 4.**
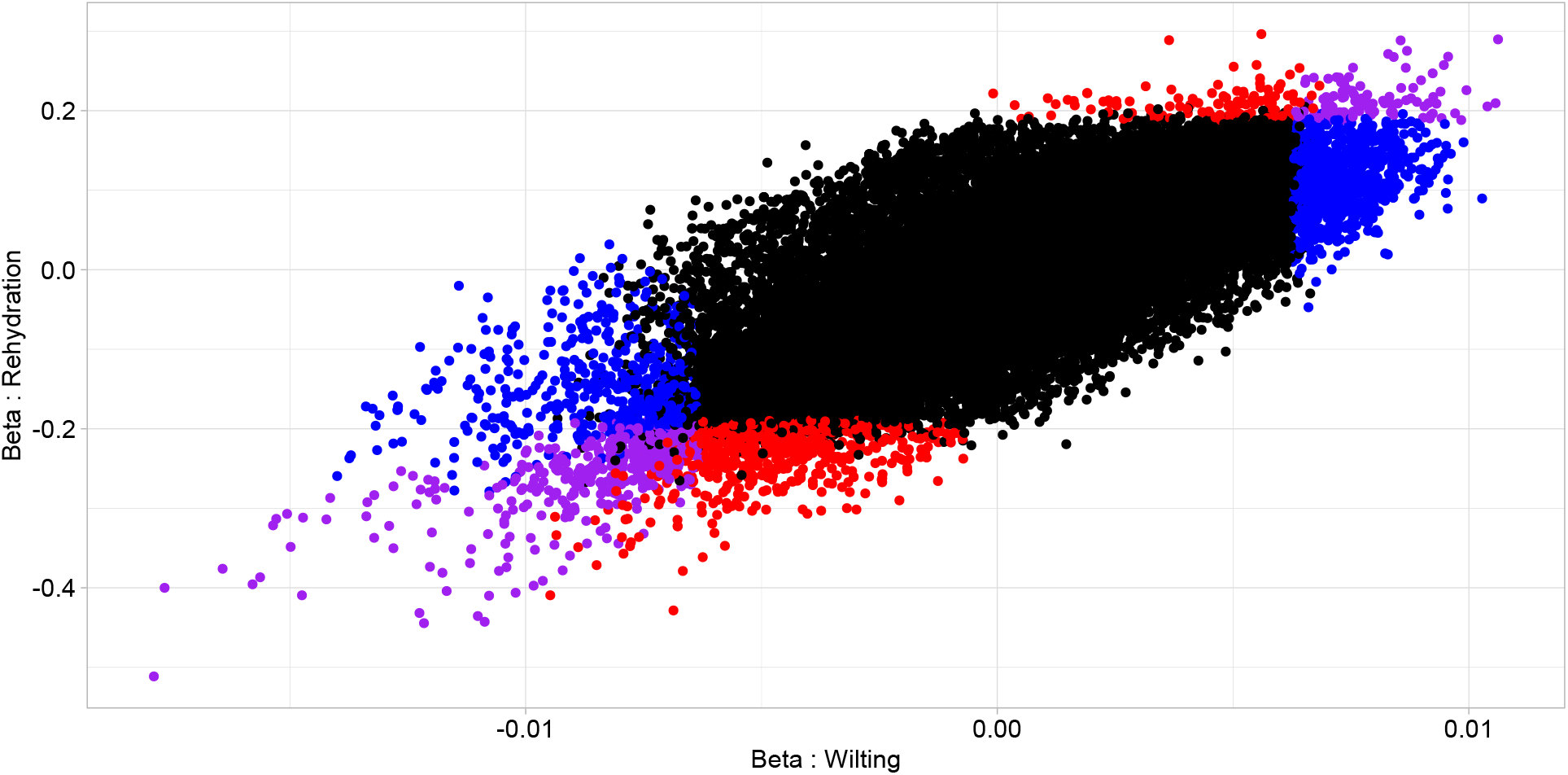
Correlations between model beta estimates from wilt VWC and wilt survival associations with RNAseq. Each point represents estimates for one transcript. Transcripts with purple dots have significant association of expression with both phenotypes, blue dots have significant association of expression with wilt VWC, red dots with wilt survival and black dots with neither phenotype.

Genes with expression strongly associated with wilt survival had functions more explicitly associated with detoxifying reactive oxygen species and negative products of photorespiration as well as continued light harvesting. The gene with the strongest association with wilt survival was a homolog of *DegP5* – a gene that dampens Ca^+2^ signaling associated with stress responses and is associated with reduced ROS (Fig. 5A). This gene also had the strongest positive association with wilt VWC of any gene, that is, high expression of this gene is linked with wilting at a high VWC. Expression of a *CUT1* (*Cutinase 1*) homolog, a gene that impacts cuticle wax biosynthesis, was also strongly associated with greater survival following wilting. One additional candidate gene, a homolog of *SAPK2*, also had among the highest associations with increased wilt survival and higher wilt VWC (Fig. 5C). Other genes, including homologs of *PGLP2*, *CAB7*, and *LHCA6*, with expression strongly associated with wilt survival are more weakly associated with wilt VWC. These genes have less clear associations with known drought responses. Together, these candidates suggest that genes associated with lower wilt VWC include those typically associated with dehydration avoidance, while genes associated with wilt survival are associated with increasing photosynthesis and protecting tissues against the harmful bioproducts of stress responses.

**Fig. 5.**
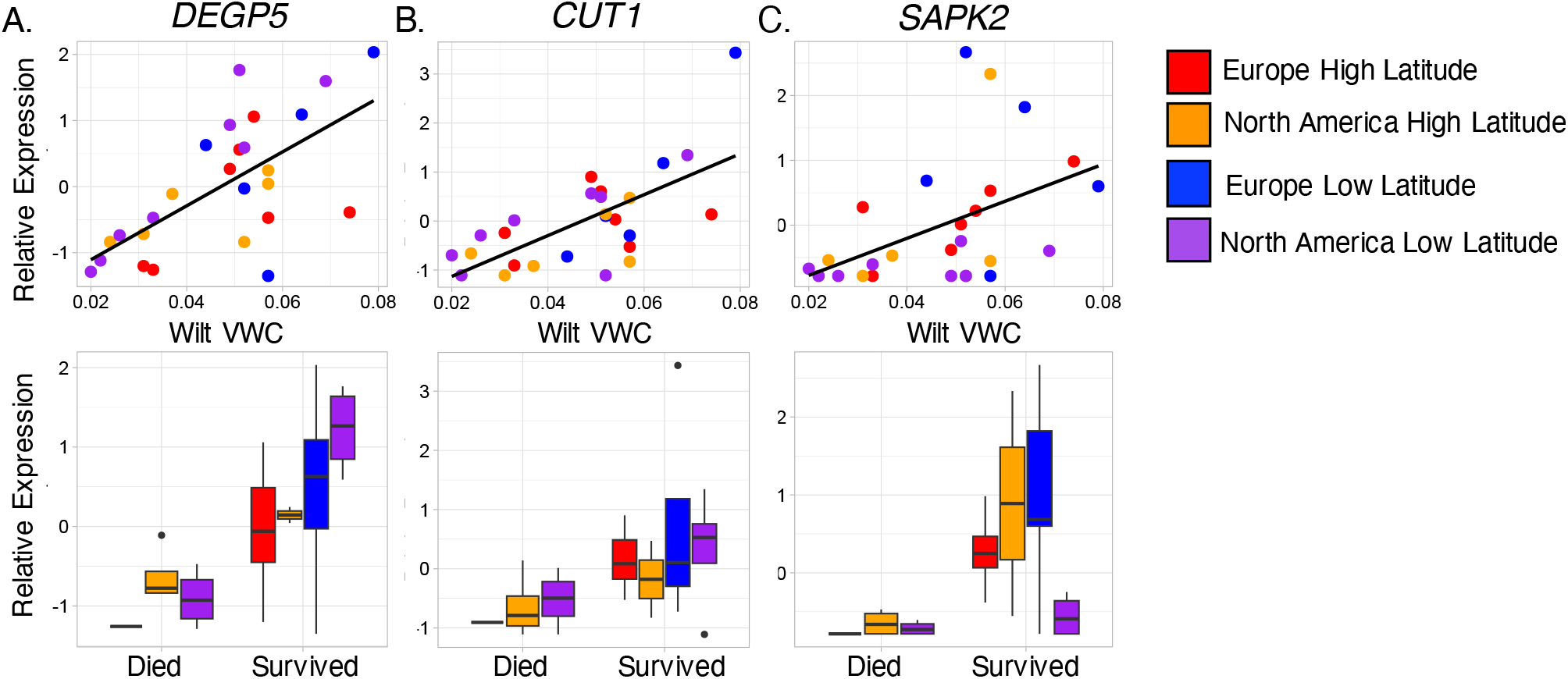
Candidate genes associated with wilt survival. Lower wilt VWC indicates greater dehydration avoidance while higher wilt survival indicates greater dehydration tolerance. Points and boxplots are colored by latitude and range: low latitude Europe = blue, high latitude Europe = red, low latitude North America = purple, High Latitude North American = orange. Each point represents a different individual. Gene Abbreviations: *DeGP5* = degradation of periplasmic proteins 5; *CUT1* = Cutinase 1; *SAPK2* = sucrose non-fermenting 1-related kinase 2.

### Expression patterns associated with phenotypic differences between ranges

While many genes are associated with dehydration avoidance and tolerance phenotypes, a limited subset of these candidates have the potential to drive the differences in dehydration avoidance or tolerance between ranges. We examined which of the candidate genes linked to dehydration avoidance or tolerance also had differential expression across ranges or latitudes. As is the case with many genes in the turquoise network, relative expression of *BCAT2* in the drought treatment is greater in high latitude populations, exhibits variable expression in low latitude North American populations, and has very limited expression in the low latitude European populations (Range:Latitude: log2FoldChange = 30, P = 4.83e-9; Fig. 3, Fig. S8).

Similar patterns exist for both *P5CS* (Range:Latitude: log2FoldChange = 29.9, P = 8.28e-11) and *GOLS2* (Range:Latitude: log2FoldChange = 18.5, P = 0.002). To determine other genes that may create phenotypic differences between ranges or across latitudes, we identified genes whose relative expression is best explained by an interaction between wilt VWC or wilt survival and range. Many genes (22.8% of genes surveyed; Table S8) exhibited such interactions including 46 of the 52 genes in the turquoise network. These results suggest the molecular variation underlying phenotypic differences between ranges either involves many loci of varying effect sizes or are large effect variants that impact substantial hormonal stress responses and many gene expression networks.

## DISCUSSION

During invasions, plants encounter novel environments and may need to acclimate or adapt to succeed. Here we identify differences in drought resistance strategies that have evolved following the introduction of white clover to North America from its native range in Europe. North American populations have stronger dehydration avoidance responses while European populations have stronger dehydration tolerance responses. These differences appear to be shaped by the abiotic conditions in each region as well as a strong tradeoff between dehydration avoidance and tolerance. We also identify a number of strong candidate genes underlying each of these phenotypes with a significant proportion falling into a single co-expression network. Most of these genes have expression profiles where higher expression improves either dehydration avoidance or tolerance while decreasing the other phenotype. Additionally, the limited subset of genes that are associated with both a drought resistant phenotype and exhibit differential expression between native and introduced ranges represent the best candidates for identifying a molecular basis of drought adaptation following introduction. We discuss these results below in the context of invasion biology and the evolutionary genetics of adaptation to plant abiotic stresses: two fields that overlap but are rarely synthesized.

### Evolution of Drought Resistance During Invasions

The roles of selection and adaptation during invasions is now widely documented with many species, including white clover, rapidly evolving climate-associated clines following introduction (e.g. Kooyers & Olsen, 2012; Colautti & Barrett, 2013; Battlay *et al*., 2023). However, generalization of which traits are favored during invasions is less clear (Kolar and Lodge 2001). Different trait-based hypotheses suggest that rapid growth (EICA; Blossey & Notzold, 1995) or stress tolerance may be favored in the introduced regions (Gewing *et al*. 2019). Our results are partially consistent with both hypotheses, but are far more nuanced as drought resistance is conferred through many different mechanisms. Introduced North American populations wilt at lower volumetric water contents than populations from the native European range, but also are less able to tolerate wilting than native populations (Fig. 1). Thus, North American populations are likely more dehydration avoidant than European populations. Greater dehydration avoidance may allow plants to maintain growth during periods of low water and high evapotranspiration.

This would be consistent with devoting more resources to rapid growth within the introduced range as well as being more resistant to initial water stress. This nuance may account for some of the mixed evidence for the EICA within the literature (Rotter and Holeski 2018).

We suggest that the evolution of greater stress avoidance to facilitate rapid growth may be a pattern frequently observed across invasions of perennial plants. Introduced species are likely to encounter different abiotic regimes in their new range. Invasive populations of annual plants often evolve more rapid life histories (Colautti and Barrett 2013; Battlay *et al*. 2023), but perennial plants must survive stresses in order to establish and reproduce. There are few plant systems where the evolution of drought avoidance or tolerance have been assessed across native and introduced regions. These examples largely consist of species where it is clear that they have a very different drought resistance strategy in the introduced range. For instance, camphorweed (*Heteroteca subaxcillaris*) has drastically greater root growth in its invasive range in Israel than in its native range, North America, that likely is a benefit both for low water availability and structural integrity in extremely arid dune environments in Israel (Sternberg 2016). Jerusalem artichokes (*Helianthus tuberous*) have evolved extreme clonality phenotypes in their introduced range (Europe) to accommodate a shift to a more aquatic environment (Bock *et al*. 2018).

Outside of plants, a greater tolerance or plasticity to stressful environments is often observed – for instance, anoles (*Anolis cristatellus)* from the introduced range survive colder temperatures than from the native range (Leal and Gunderson 2012). We suggest that invasive species should be more widely used for identifying natural phenotypic and genomic variation in drought resistance.

While our results are consistent with classic invasion biology theory, they also make sense in the context of environmental characteristics of the populations. The populations involved in this experiment have very different growing seasons and historical climatic features (Table S1).

Southern European populations come from a Mediterranean climate with cool and temperate winters and hot dry summers. Relative to other locations, there is little precipitation throughout the year and almost no rainfall during summer months. A long summer with little rain is the ideal conditions for the evolution of dehydration tolerance in a perennial plant that likely needs to survive across multiple growing seasons (Kooyers 2015). Alternatively, eastern North America has more substantial rainfall throughout the growing season with drought stress likely coming during shorter stretches without rain (Table S1). These conditions may favor the evolution of an avoidance strategy as additional rainfall is likely coming in the near future. Interestingly, we observe very little difference in drought resistance strategy between high and low latitude populations of white clover in the native or introduced ranges. In both ranges, on average, lower latitude populations are less dehydration avoidant but more dehydration tolerant. This is surprising as previous work documents differentiation in life history strategy evolving across this gradient with warmer southern populations employing a more annualized life history (Wright *et al*. 2021). We suggest our small sample size only has the power to detect major effects and future larger studies may be able to parse differences in stress responses across latitudinal gradients.

### Molecular basis of drought tolerance and avoidance variation

The molecular basis of drought resistance has been examined extensively in crop plants and model systems, but there has been limited success in determining the genes that underlie variation in natural populations (Des Marais *et al*. 2013; VanWallendael *et al*. 2019). Here we find several candidate genes that likely influence dehydration avoidance and tolerance responses across both invasive and native ranges despite relatively few mapped reads from only a single tissue type (Fig. 3). Notably the turquoise co-expression network is strongly enriched for gene ontologies related to drought responses (i.e. response to abiotic stress, response to water and proline synthesis). A high proportion of the genes within this network are also associated with wilt VWC including some of the strongest candidate genes. Interestingly, these genes represent a diverse set of drought-related functions in other organisms.

*NAC4* and *SDIR* homologs are likely involved in stress signaling pathways. NAC4 (nascent polypeptide-associated complex) is a transcription factor that is strongly upregulated in response to drought, salt, and water stress in many species (Xia *et al*. 2010; Shao *et al*. 2015; Yu *et al*. 2016) and confers drought tolerance when transgenically expressed in *Arabidopsis* (Mei *et al*. 2021). More generally, the NAC gene family is involved in stomatal closure and in lateral root development in rice (Hu *et al*. 2006; Zheng *et al*. 2009). SDIR1 (Salt and Drought-Induced Ring Finger1) is a RING-finger E3 ligase that is upregulated in *Arabidopsis* in response to drought stress and modulates both the ABA response and stomatal closure (Zhang *et al*. 2007). While upregulation of both genes in response to drought stress in our system is consistent with findings in other species and expected for dehydration avoidant plants, *sdir1* mutants in *Arabidopsis* had greater mortality following wilting than wild type plants. This suggests an additional dehydration tolerance role for SDIR1 that we do not observe.

Expression of multiple genes that promote the synthesis of osmoprotectants (i.e. branched chain amino acids or BCAAs, proline, and raffinose family oligosaccharides) are also strongly associated with lower wilt VWC. *BCAT2* (BCAA transaminase 2) catalyzes the final transamination step in production of BCAAs as well as the first degradation step. Under stress, BCAAs accumulate as osmoprotectants in an ABA-dependent manner, with BCAT serving as the main regulator in *Arabidopsis*. P5CS, delta 1-pyrroline-5-carboxylate synthase, controls the rate-limiting step of proline biosynthesis from glutamate during osmotic and salt stress (Strizhov *et al*. 1997). Transgenic overexpression of P5CS consistently increases tolerance to drought and salt stress across several plant species by increasing proline content (Su and Wu 2004; Yamchi *et al*. 2007; Vendruscolo *et al*. 2007). *GOLS2* (galactinol synthase 2) encodes galactinol synthase 2, a key enzyme that regulates the first step in biosynthesis of raffinose family oligosaccharides resulting in the creation of galactinol (Panikulangara *et al*. 2004). While the oligosaccharides act as osmoprotectants and as antioxidants, both galactinol and the oligosaccharides may also act as signaling molecules (Song *et al*. 2016). In other species, GOLS activity is triggered by abiotic and biotic stress (*Arabidopsis*, Coffee, Rice) and confers drought tolerance (Dos Santos and Vieira 2020). Unexpectedly, our results indicate that expression of *GOLS2* is linked with lower wilt VWC (greater dehydration avoidance) and with lower survival following wilting (lower dehydration tolerance). Increased expression of *GOLS2* in *Arabidopsis* does reduce transpiration rate (Taji *et al*. 2002).

We also note a third process that may enhance dehydration avoidance and tolerance responses: the recycling and efficient use of nitrogen. *AlaAT2* (alanine aminotransferase 2) is strongly upregulated in plants that wilt at low VWC. We propose that this upregulation links drought response to nitrogen metabolism. Specifically, AlaAT2 catalyzes the reversible conversion of alanine and α-oxoglutarate to pyruvate and glutamate, facilitating the synthesis of other amino acids as needed (Miyashita *et al*. 2007). *AlaAT* homologs are upregulated during hypoxia in several species, including *Arabidopsis*, Poplar, Medicago, and Corn, and are regulated by ABA in other stressful contexts (Ricoult *et al*. 2005; Miyashita *et al*. 2007; Xu *et al*. 2017). Drought stress typically decreases the activity of nitrate reductase and nitrile reductase (Foyer *et al*. 1998), impairing the plant’s ability to accumulate amino acids due to limited nitrogen resources. Greater AlaAT2 activity could convert alanine into glutamate, which may then be used to produce BCAAs or proline as osmoprotectants. In sum, we hypothesize that the production of osmoprotectants requires carbon and nitrogen resources that may be most easily procured through recycling of existing metabolites.

The candidate genes associated with wilt survival generally do not have strong functional ties to drought responses in other systems with two exceptions. *DegP5* (degradation of periplasmic proteins 5) belongs to a family of proteases that is typically upregulated in response to abiotic stress. Overexpression of *DegP5* has been linked to increased dehydration tolerance in *Nicotiana tabacum* via the suppression of Ca^+2^ and flagellin signaling leading to a reduced accumulation of reactive oxygen species (Chen *et al*. 2023). In our study, increased expression of *DegP5* is also associated with wilting at higher soil moistures. This suggests that the increase wilt survival comes at a cost of a less active drought response. The second exception is *SAPK2*, a plant-specific protein kinase in the family of SNF1-related protein kinase 2 (*SnRK2*) genes. Expression of the *SAPK2* homolog is positively associated with wilt survival (Fig. 5C). This role is similar to that documented in rice, where SAPK2 not only upregulates genes involved in osmoprotection but also induces genes involved in enzymatic ROS detoxification (Lou *et al*. 2017). However, a role in dehydration avoidance has not been documented. Other SnRK2s do regulate cation channels (KAT1) and impact stomatal closure in *Arabidopsis* (Sato *et al*. 2009). Other genes strongly associated with high wilt survival in our study function in other organisms in the detoxification of photorespiration byproducts (*PGLP2*; Schwarte & Bauwe, 2007) and in efficient light harvesting and photosynthesis (*LHCA6*, *CAB7*; Otani *et al*., 2017). Together, these results suggest that improved survival after wilting is achieved through a combination of dampening responses to drought, protecting cells from harmful bioproducts of drought stress, and maintaining metabolic functions.

### A tradeoff between drought avoidance and tolerance strategies?

Drought resistance strategies are often presented as non-mutually exclusive with the recognition that they represent different physiological mechanisms for a plant to acclimate to low water availability. The relationship between drought escape and dehydration avoidance has been the most extensively scrutinized as there are clear physiological tradeoffs for C3 plants – closing stomata for better dehydration avoidance leads to lower carbon uptake and the inability to synthesize more sugars for rapid growth and reproduction (Geber and Dawson 1990; Dudley 1996). Theoretical and empirical identification of tradeoffs between dehydration avoidance and tolerance is rare – potentially because there is not a true physiological tradeoff that exists or because these strategies may be expressed sequentially as water stress increases (Volaire 2018).

We document relatively strong tradeoffs between wilt VWC and wilt survival (Fig. 2) and expression responses for wilt VWC and wilt survival are strongly correlated (Fig. 4). However, the functions of genes associated with each of these phenotypes suggest that our experiment is not truly measuring separate dehydration avoidance and tolerance responses. Rather, our wilt VWC treatment is likely catching both dehydration avoidance and tolerance responses – plants that survive to lower wilting *are* effectively tolerating dehydration for longer than plants that wilted earlier, likely by increasing osmoprotectants as well as closing stomata to regulate gas exchange. Correlated dehydration avoidance and tolerance responses make sense as both functions are at least partly regulated by ABA. However, since dehydration tolerance responses often involve the production of various osmoprotectants and closing stomata limits CO2 (and useable plant nitrogen) uptake, correlated physiological responses must require recycling carbon and nitrogen.

The strong correlation between wilt VWC and wilt survival more likely represents a tradeoff between different elements of dehydration tolerance. Wilting occurs when the turgor pressure within cells falls to zero, leading to potential cellular damage. This damage is exacerbated under prolonged drought stress, as increased ROS production can overwhelm the plant’s antioxidant defenses and, coupled with stomatal closure, can also increase photorespiration flux (Noctor *et al*. 2002). To survive wilting, plants must be able to protect cellular function and integrity. Our study found that plants with better wilting survival had higher expression of genes associated with antioxidant defense (*DegP5*) and detoxification of photorespiration byproducts (*PGLP2*). While expression of each of these genes was associated with high wilt VWC, the only truly strong correlation was for *DegP5* – a gene linked to diminishing Ca^+2^ signaling as well as lower ROS production in tobacco (Chen *et al*. 2023). Expression of this gene may represent a true tradeoff between different physiological strategies. Generally, our results suggest that a dehydration tolerance strategy actually represents multiple different types of tolerance – one strategy that provides resistance against dehydration and another that allows longer-term survival following wilting (i.e. extreme dehydration). We hypothesize that dehydration tolerance related to drought survival may have overlapping physiology with desiccation tolerance strategies occurring in plants like resurrection fern. In sum, connecting variation between physiological processes and plant performance will provide additional insight into the evolution of different ecological strategies in specific environments.

### Loci underlying drought response adaptation following introduction

Identifying the causative molecular variants that underlie natural intraspecific variation in drought responses has proven challenging with limited of successes (Juenger 2013; VanWallendael *et al*. 2019). This objective is largely beyond the scope of our study. We do identify many genes that are both linked with wilt VWC or wilt survival phenotypes and also show variable expression across ranges, conditions and latitudes. This overlap includes a substantial portion of the turquoise gene co-expression network, suggesting that variation in dehydration avoidance and tolerance phenotypes is physiological manifested through broad shifts across gene expression networks (e.g. Lovell *et al*., 2018) rather than through simple, large effect variants that directly cause lower stomatal conductance or greater osmoprotection (e.g. Kesari *et al*., 2012; Des Marais *et al*., 2014).

Recent work has identified several haploblocks that underlie climate adaptation following introduction to North America as well as several candidate genes within haploblocks that were differentially expressed across drought treatment, range and latitude (Battlay *et al*. 2024). Within the three haploblocks on Chr04_Occ (syn. Chromosome 7), 26 differentially expressed transcripts were found (9 on hb7a1; 14 on hb7a2; and 3 hb7b, Table S9). On hb7a1, 4 genes within the turquoise submodule were differentially expressed. Two turquoise submodule genes, *P5CS1 and FER1*, were strongly correlated with wilt VWC and wilt survival. In addition, *BRR2A* and *DWF5* were correlated significantly with wilt survival whereas *IAA6* and *DWF5* were correlated significantly with wilt VWC. On hb7b, transcripts belonging to the gene *CCL12* are significantly correlated with wilting. Multiple of these genes have allele variation associated with higher fitness in North American common gardens (Albano et al. in prep; Battlay *et al*., 2024). These results are a first step to tying locally adaptive genetic variation to variation in gene regulatory networks, physiology and ecological strategies.

### Conclusions

Our work suggests that selection acts on rapid timescales following introduction to allow more precise adaptation of drought responses to divergent environmental conditions. Specifically, selection favors specific drought resistance strategies, with tradeoffs occurring between strategies at both phenotypic and molecular levels. We identify a co-expression network that underlies both dehydration avoidance and some elements of dehydration tolerance where many of these loci also exhibit differential expression patterns between the native and introduced ranges of white clover. We also identify tradeoffs at the molecular level among functions (osmoprotection and detoxification) typically identified within the dehydration tolerance strategy – a result that needs to be carefully considered in breeding efforts. This work stimulates more studies examining and relating genetic variation and expression networks to phenotypic differences within natural populations to create a broader consensus on mechanisms creating variation in drought resistance and survival.

## ACKNOWLEDGEMENTS

The Louisiana Optical Network (LONI) provided computing resources for RNAseq analysis. We thank Simon Innes and Marc Johnson for seed collections. We thank Happy Gill, Jashandeep Nijjar, and Ruth Thomas for assistance developing germplasm. This work was funded through an NSF grant to NJK (OIA-1920858). Seed collections and propagation was funded by a Canada Research Chair (950-231981), EWR Steacie Fellowship (544292), NSERC Discovery Grant (RGPIN-2016-06063), and NSERC RTI Grant (EQPEQ 423691) to Marc Johnson as well as a NSERC CGS Doctoral Award to LJA.

## DATA AVAILABILITY

Phenotypic data from the growth chamber experiment is available via dryad: link. RNAseq data is available via NCBI’s short read archive: link.

## .SUPPLEMENTAL INFORMATION

Table S1. Geographic and environmental characteristics of white clover populations.

Table S2. Mapping statistics from RNAseq experiment

Table S3. Univariate model results examining associations between drought strategy phenotypes across ranges, latitudes, and treatments.

Table S4. Univariate model results examining associations between drought strategy phenotypes with water-availability related variables of collection populations

Table S5. Summary of differentially expressed genes with associated modules and phenotypic associations.

Table S6. Gene Ontology analysis of WGCNA modules

Table S7. Genes with variation in expression associated with wilt VWC or wilt survival

Table S8. Genes with significant range:phenotype interactions.

Table S9. Patterns of expression within haploblocks

Fig. S1. Map of sampling locations.

Fig. S2. Volumetric water content of pots within well-watered and dry down treatments throughout the experiment.

Fig. S3. Relationships between drought strategy phenotypes and leaf number at the start of the experiment.

Fig. S4. Associations between water-availability related variables of collection populations and leaf number at the start of the experiment

Fig. S5. Associations between water-availability related variables of collection populations and drought avoidance and tolerance.

Fig. S6. Venn diagram of differentially expressed genes across latitude, range and treatment contrasts.

Fig. S7. Normalization expression profiles across samples.

Fig. S8. Relative expression across treatment, region, and latitude for candidate genes

## REFERENCES

1. Altschul SF, Gish W, Miller W, Myers EW, Lipman DJ. 1990. Basic local alignment search tool. Journal of Molecular Biology 215: 403–410.

2. Ashburner M, Ball CA, Blake JA, et al. 2000. Gene Ontology: tool for the unification of biology. Nature Genetics 25: 25–29.

3. Aslam MM, Waseem M, Jakada BH, et al. 2022. Mechanisms of Abscisic Acid-Mediated Drought Stress Responses in Plants. International Journal of Molecular Sciences 23: 1084.

4. Barabási A-L, Albert R. 1999. Emergence of scaling in random networks. Science 286: 509– 512.

5. Battlay P, Hendrickson BT, Mendez-Reneau JI, et al. 2024. Structural variants underlie parallel adaptation following global invasion. bioRxiv. doi: 10.1101/2024.07.09.602765

6. Battlay P, Wilson J, Bieker VC, et al. 2023. Large haploblocks underlie rapid adaptation in the invasive weed Ambrosia artemisiifolia. Nature Communications 14: 1717.

7. Belda-Palazón B, Adamo M, Valerio C, et al. 2020. A dual function of SnRK2 kinases in the regulation of SnRK1 and plant growth. Nature Plants 6: 1345–1353.

8. Blossey B, Notzold R. 1995. Evolution of increased competitive ability in invasive nonindigenous plants: A hypothesis. The Journal of Ecology 83: 887.

9. Bock DG, Kantar MB, Caseys C, Matthey-Doret R, Rieseberg LH. 2018. Evolution of invasiveness by genetic accommodation. Nature Ecology & Evolution 2: 991–999.

10. Bonnamour A, Gippet JMW, Bertelsmeier C. 2021. Insect and plant invasions follow two waves of globalisation (J Gurevitch, Ed.). Ecology Letters 24: 2418–2426.

11. Bouzid M, He F, Schmitz G, et al. 2019. *Arabidopsis* species deploy distinct strategies to cope with drought stress. Annals of Botany XX: 1–14.

12. Chen G, Shu Y, Jian Z, Duan L, Mo Z, Liu R. 2023. The NtDEGP5 gene improves drought tolerance in tobacco ( *Nicotiana tabacum* L.) by dampening plastid extracellular Ca2+ and flagellin signaling and thereby reducing ROS production. Plant Molecular Biology 113: 265– 278.

13. Chen S, Zhou Y, Chen Y, Gu J. 2018. fastp: an ultra-fast all-in-one FASTQ preprocessor. Bioinformatics 34: i884–i890.

14. Colautti RI, Barrett SCH. 2013. Rapid adaptation to climate facilitates range expansion of an invasive plant. Science 342: 364–366.

15. Colautti RI, Lau JA. 2015. Contemporary evolution during invasion: evidence for differentiation, natural selection, and local adaptation. Molecular Ecology 24: 1999–2017.

16. Dar NA, Amin I, Wani W, et al. 2017. Abscisic acid: A key regulator of abiotic stress tolerance in plants. Plant Gene 11: 106–111.

17. Des Marais DL, Auchincloss LC, Sukamtoh E, et al. 2014. Variation in *MPK12* affects water use efficiency in Arabidopsis and reveals a pleiotropic link between guard cell size and ABA response. Proceedings of the National Academy of Sciences 111: 2836–2841.

18. Des Marais DL, Hernandez KM, Juenger TE. 2013. Genotype-by-environment interaction and plasticity: Exploring genomic responses of plants to the abiotic environment. Annual Review of Ecology, Evolution, and Systematics 44: 5–29.

19. Dos Santos TB, Vieira LGE. 2020. Involvement of the galactinol synthase gene in abiotic and biotic stress responses: A review on current knowledge. Plant Gene 24: 100258.

20. Dudley SA. 1996. Differing selection on plant physiological traits in response to environmental water availibility: A test of adaptive hypotheses. Evolution 50: 92–102.

21. Fick SE, Hijmans RJ. 2017. WorldClim 2: new 1-km spatial resolution climate surfaces for global land areas. International Journal of Climatology 37: 4302–4315.

22. FitzPatrick JA, Doucet BI, Holt SD, Patterson CM, Kooyers NJ. 2023. Unique drought resistance strategies occur among monkeyflower populations spanning an aridity gradient. American Journal of Botany 110: e16207.

23. Fox J, Friendly M, Weisberg S. 2013. Hypothesis tests for multivariate linear models using the car package. R Journal 5: 39–52.

24. Foyer CH, Valadier M-H, Migge A, Becker TW. 1998. Drought-induced effects on Nitrate Reductase activity and mRNA and on the coordination of nitrogen and carbon metabolism in Maize leaves. Plant Physiology 117: 283–292.

25. Geber MA, Dawson TE. 1990. Genetic variation in and covariation between leaf gas exchange, morphology, and development in *Polygonum arenastrum*, an annual plant. Oecologia 85: 153– 158.

26. Gewing M-T, Goldstein E, Buba Y, Shenkar N. 2019. Temperature resilience facilitates invasion success of the solitary ascidian *Herdmania momus*. Biological Invasions 21: 349–361.

27. Hsu P, Dubeaux G, Takahashi Y, Schroeder JI. 2021. Signaling mechanisms in abscisic acid- mediated stomatal closure. The Plant Journal 105: 307–321.

28. Hu H, Dai M, Yao J, et al. 2006. Overexpressing a NAM, ATAF, and CUC (NAC) transcription factor enhances drought resistance and salt tolerance in rice. Proceedings of the National Academy of Sciences 103: 12987–12992.

29. Huey RB. 2000. Rapid evolution of a geographic cline in size in an introduced fly. Science 287: 308–309.

30. Innes SG, Santangelo JS, Kooyers NJ, Olsen KM, Johnson MTJ. 2022. Evolution in response to climate in the native and introduced ranges of a globally distributed plant. Evolution 76: 1495–1511.

31. Jahufer MZZ, Cooper M, Ayres JF, Bray RA. 2002. Identification of research to improve the efficiency of breeding strategies for white clover in Australia - a review. Australian Journal of Agricultural Research 53: 239.

32. Juenger TE. 2013. Natural variation and genetic constraints on drought tolerance. Current Opinion in Plant Biology 16: 274–281.

33. Kanehisa M, Goto S. 2000. KEGG: Kyoto Encyclopedia of Genes and Genomes. Nucleic Acids Research 28: 27–30.

34. Kesari R, Lasky JR, Villamor JG, et al. 2012. Intron-mediated alternative splicing of *Arabidopsis P5CS1* and its association with natural variation in proline and climate adaptation. Proceedings of the National Academy of Sciences 109: 9197–9202.

35. Kjærgaard T. 2003. A plant that changed the world: The rise and fall of clover 1000-2000. Landscape Research 28: 41–49.

36. Kolar CS, Lodge DM. 2001. Progress in invasion biology: predicting invaders. Trends in Ecology & Evolution 16: 199–204.

37. Kooyers NJ. 2015. The evolution of drought escape and avoidance in natural herbaceous populations. Plant Science 234: 155–162.

38. Kooyers NJ, Gage LR, Al-Lozi A, Olsen KM. 2014. Aridity shapes cyanogenesis cline evolution in white clover (*Trifolium repens* L.). Molecular Ecology 23: 1053–1070.

39. Kooyers NJ, Greenlee AB, Colicchio JM, Oh M, Blackman BK. 2015. Replicate altitudinal clines reveal that evolutionary flexibility underlies adaptation to drought stress in annual *Mimulus guttatus*. New Phytologist 206: 152–165.

40. Kooyers NJ, Olsen KM. 2012. Rapid evolution of an adaptive cyanogenesis cline in introduced North American white clover (*Trifolium repens* L.). Molecular Ecology 21: 2455–2468.

41. Kooyers NJ, Olsen KM. 2013. Searching for the bull’s eye: Agents and targets of selection vary among geographically disparate cyanogenesis clines in white clover (*Trifolium repens* L.). Heredity 111: 495–504.

42. Langfelder P, Horvath S. 2008. WGCNA: an R package for weighted correlation network analysis. BMC Bioinformatics 9: 559.

43. Langmead B, Salzberg SL. 2012. Fast gapped-read alignment with Bowtie 2. Nature Methods 9: 357–360.

44. Leal M, Gunderson AR. 2012. Rapid change in the thermal tolerance of a tropical lizard. The American Naturalist 180: 815–822.

45. Lou D, Wang H, Liang G, Yu D. 2017. OsSAPK2 Confers Abscisic Acid Sensitivity and Tolerance to Drought Stress in Rice. Frontiers in Plant Science 8: 993.

46. Love MI, Huber W, Anders S. 2014. Moderated estimation of fold change and dispersion for RNA-seq data with DESeq2. Genome Biology 15.

47. Lovell JT, Jenkins J, Lowry DB, et al. 2018. The genomic landscape of molecular responses to natural drought stress in *Panicum hallii*. Nature Communications 9: 5213.

48. Ludlow MM. 1989. Strategies of response to water stress In: Structural and Functional Responses to Environmental Stresses. The Hague: SPB Academic Publishing, 269–281.

49. Lunn JE, Delorge I, Figueroa CM, Van Dijck P, Stitt M. 2014. Trehalose metabolism in plants. The Plant Journal 79: 544–567.

50. Maron JL, Vilà M, Bommarco R, Elmendorf S, Beardsley P. 2004. Rapid evolution of an invasive plant. Ecological Monographs 74: 261–280.

51. Marshall AH, Rascle C, Abberton MT, Michaelson-Yeates TPT, Rhodes I. 2001. Introgression as a route to improved drought tolerance in white clover (*Trifolium repens* L.). Journal of Agronomy and Crop Science 187: 11–18.

52. McKay JK, Richards JH, Mitchell-Olds T. 2003. Genetics of drought adaptation in *Arabidopsis thaliana* I. Pleiotropy contributes to genetic correlations among ecological traits. Molecular Ecology 12: 1137–1151.

53. Mei F, Chen B, Li F, et al. 2021. Overexpression of the wheat NAC transcription factor TaSNAC4-3A gene confers drought tolerance in transgenic Arabidopsis. Plant Physiology and Biochemistry 160: 37–50.

54. Miyashita Y, Dolferus R, Ismond KP, Good AG. 2007. Alanine aminotransferase catalyses the breakdown of alanine after hypoxia in *Arabidopsis thaliana*. The Plant Journal 49: 1108–1121.

55. Noctor G, Veljovic-Jovanovic S, Novitskaya L, Foyer CH. 2002. Drought and oxidative load in the leaves of C3 plants: a predominant role for photorespiration? Annals of Botany 89: 841– 850.

56. Otani T, Yamamoto H, Shikanai T. 2017. Stromal loop of Lhca6 is responsible for the linker function required for the NDH–PSI supercomplex formation. Plant and Cell Physiology 58: 851–861.

57. Panikulangara TJ, Eggers-Schumacher G, Wunderlich M, Stransky H, Schöffl F. 2004. *Galactinol synthase1* . A novel heat shock factor target gene responsible for heat-induced synthesis of raffinose family oligosaccharides in Arabidopsis. Plant Physiology 136: 3148–3158.

58. Patro R, Duggal G, Love MI, Irizarry RA, Kingsford C. 2017. Salmon provides fast and bias- aware quantification of transcript expression. Nature Methods 14: 417–419.

59. Pertea G, Pertea M. 2020. GFF Utilities: GffRead and GffCompare. F1000Research 9: 304.

60. Pyšek P, Richardson DM. 2007. Traits associated with invasiveness in alien plants: Where do we stand? In: Nentwig W, ed. Ecological Studies. Biological Invasions. Berlin, Heidelberg: Springer Berlin Heidelberg, 97–125.

61. Raghavendra AS, Gonugunta VK, Christmann A, Grill E. 2010. ABA perception and signalling. Trends in Plant Science 15: 395–401.

62. Richards CL, Bossdorf O, Muth NZ, Gurevitch J, Pigliucci M. 2006. Jack of all trades, master of some? On the role of phenotypic plasticity in plant invasions. Ecology Letters 9: 981– 993.

63. Ricoult C, Cliquet J, Limami AM. 2005. Stimulation of alanine amino transferase ( *AlaAT* ) gene expression and alanine accumulation in embryo axis of the model legume *Medicago truncatula* contribute to anoxia stress tolerance. Physiologia Plantarum 123: 30–39.

64. Rotter MC, Holeski LM. 2018. A meta-analysis of the evolution of increased competitive ability hypothesis: genetic-based trait variation and herbivory resistance trade-offs. Biological Invasions 20: 2647–2660.

65. Santangelo JS, Battlay P, Hendrickson BT, et al. 2023. Haplotype-resolved, chromosome- level assembly of white clover ( *Trifolium repens* L., Fabaceae) (V Castric, Ed.). Genome Biology and Evolution 15: evad146.

66. Santangelo JS, Ness RW, Cohan B, et al. 2022. Global urban environmental change drives adaptation in white clover. Science 375: 1275–1281.

67. Sato A, Sato Y, Fukao Y, et al. 2009. Threonine at position 306 of the KAT1 potassium channel is essential for channel activity and is a target site for ABA-activated SnRK2/OST1/SnRK2.6 protein kinase. Biochemical Journal 424: 439–448.

68. Schwarte S, Bauwe H. 2007. Identification of the photorespiratory 2-Phosphoglycolate Phosphatase, *PGLP1*, in Arabidopsis. Plant Physiology 144: 1580–1586.

69. Seebens H, Blackburn TM, Dyer EE, et al. 2017. No saturation in the accumulation of alien species worldwide. Nature Communications 8: 14435.

70. Shao H, Wang H, Tang X. 2015. NAC transcription factors in plant multiple abiotic stress responses: progress and prospects. Frontiers in Plant Science 6.

71. Siepielski AM, Morrissey MB, Buoro M, et al. 2017. Precipitation drives global variation in natural selection. Science 355: 959–962.

72. Song C, Chung WS, Lim CO. 2016. Overexpression of heat shock factor gene HsfA3 Increases galactinol levels and oxidative stress tolerance in Arabidopsis. Molecules and Cells 39: 477–483.

73. Sternberg M. 2016. From America to the Holy Land: disentangling plant traits of the invasive *Heterotheca subaxillaris*. Plant Ecology 217: 1307–1314.

74. Strizhov N, Ábrahám E, Ökrész L, et al. 1997. Differential expression of two *P5CS* genes controlling proline accumulation during salt-stress requires ABA and is regulated by *ABA1, ABI1* and *AXR2* in *Arabidopsis*. The Plant Journal 12: 557–569.

75. Su J, Wu R. 2004. Stress-inducible synthesis of proline in transgenic rice confers faster growth under stress conditions than that with constitutive synthesis. Plant Science 166: 941–948.

76. Taji T, Ohsumi C, Iuchi S, et al. 2002. Important roles of drought- and cold-inducible genes for galactinol synthase in stress tolerance in *Arabidopsis thaliana*. The Plant Journal 29: 417–426.

77. Vanegas L, Rondón L, Paula G. 2024. glmtoolbox: Set of tools to data analysis using generalized linear models.

78. VanWallendael A, Soltani A, Emery NC, Peixoto MM, Olsen J, Lowry DB. 2019. A molecular view of plant local adaptation: Incorporating stress-response networks. Annual Review of Plant Biology 70: 559–583.

79. Vendruscolo ECG, Schuster I, Pileggi M, et al. 2007. Stress-induced synthesis of proline confers tolerance to water deficit in transgenic wheat. Journal of Plant Physiology 164: 1367– 1376.

80. Volaire F. 2018. A unified framework of plant adaptive strategies to drought: Crossing scales and disciplines. Global Change Biology 24: 2929–2938.

81. Wilkinson S, Davies WJ. 2010. Drought, ozone, ABA and ethylene: new insights from cell to plant to community. Plant, Cell & Environment 33: 510–525.

82. Wright SJ, Cui Zhou D, Kuhle A, Olsen KM. 2018. Continent-Wide climatic variation drives local adaptation in North American white clover. Journal of Heredity 109: 78–89.

83. Wright SJ, Goad DM, Gross BL, Muñoz PR, Olsen KM. 2021. Genetic trade-offs underlie divergent life history strategies for local adaptation in white clover. Molecular Ecology: mec.16180.

84. Wu F, Ma S, Zhou J, et al. 2021. Genetic diversity and population structure analysis in a large collection of white clover ( *Trifolium repens* L.) germplasm worldwide. PeerJ 9: e11325.

85. Xia N, Zhang G, Liu X-Y, et al. 2010. Characterization of a novel wheat NAC transcription factor gene involved in defense response against stripe rust pathogen infection and abiotic stresses. Molecular Biology Reports 37: 3703–3712.

86. Xu Z, Ma J, Qu C, et al. 2017. Identification and expression analyses of the alanine aminotransferase (AlaAT) gene family in poplar seedlings. Scientific Reports 7: 45933.

87. Yamchi A, Rastgar Jazii F, Mousavi A, Karkhane AA, Renu. 2007. Proline accumulation in transgenic tobacco as a result of expression of Arabidopsis Δ1-Pyrroline-5-carboxylate synthetase (P5CS) during osmotic stress. Journal of Plant Biochemistry and Biotechnology 16: 9–15.

88. Yoshida T, Fujita Y, Maruyama K, et al. 2015. Four *A rabidopsis* AREB / ABF transcription factors function predominantly in gene expression downstream of SNRK2 kinases in abscisic acid signalling in response to osmotic stress. *Plant*, Cell & Environment 38: 35–49.

89. Yu X, Liu Y, Wang S, et al. 2016. CarNAC4, a NAC-type chickpea transcription factor conferring enhanced drought and salt stress tolerances in Arabidopsis. Plant Cell Reports 35: 613–627.

90. Zhang Y, Yang C, Li Y, et al. 2007. SDIR1 Is a RING Finger E3 Ligase That Positively Regulates Stress-Responsive Abscisic Acid Signaling in *Arabidopsis*. The Plant Cell 19: 1912– 1929.

91. Zheng X, Chen B, Lu G, Han B. 2009. Overexpression of a NAC transcription factor enhances rice drought and salt tolerance. Biochemical and Biophysical Research Communications 379: 985–989.

92. Zomer RJ, Trabucco A, Bossio DA, Verchot LV. 2008. Climate change mitigation: A spatial analysis of global land suitability for clean development mechanism afforestation and reforestation. *Agriculture*, Ecosystems & Environment 126: 67–80.

