## Supplemental Figures for "Evolution of drought resistance strategies following the introduction of white clover (*Trifolium repens* L.)"

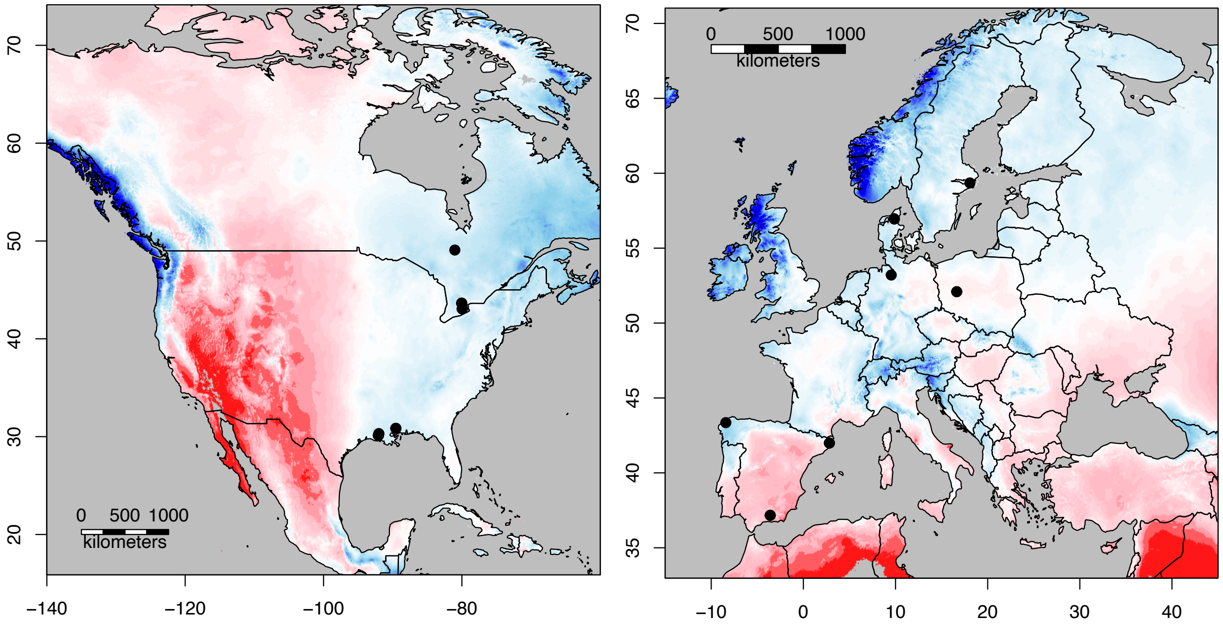


**Fig. S1.** Map of sampling locations. Black points represent sampling locations. Color raster represents annual aridity index with warmer colors indicating more xeric environments.

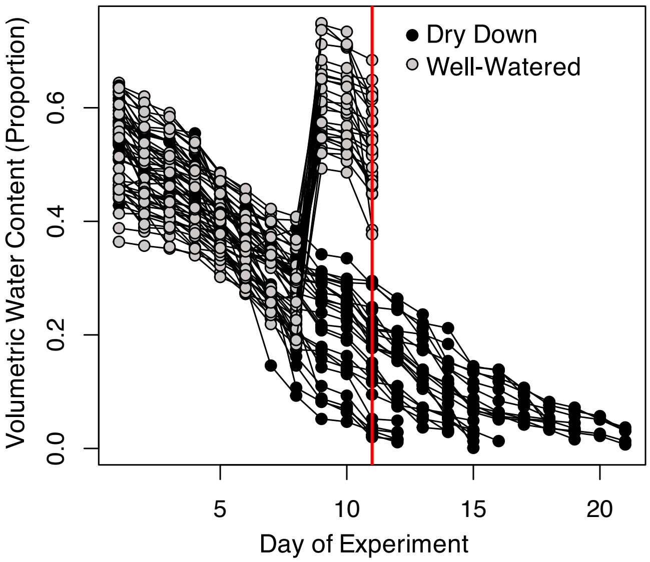


**Fig. S2** Volumetric water content of pots within well-watered (Grey) and dry down (Black) treatments throughout the experiment. Red line represents the day that tissue was taken for RNAseq analysis.

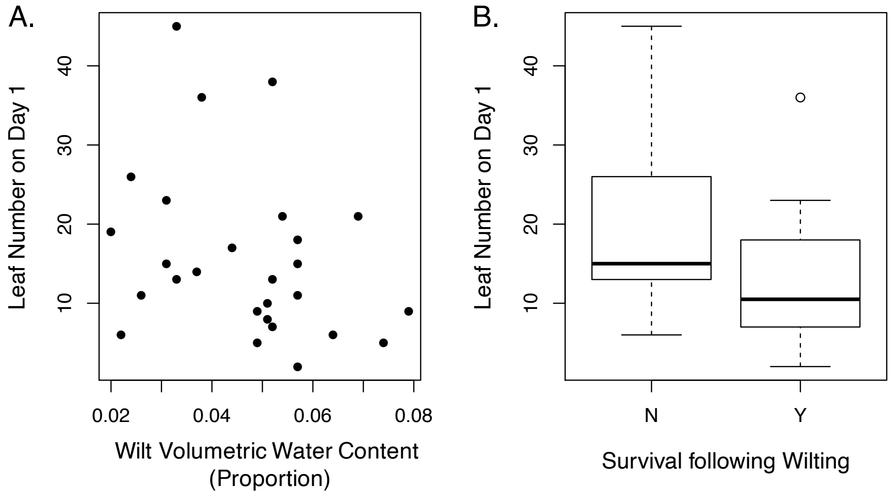


**Fig. S3** Relationships between drought strategy phenotypes and leaf number at the start of the experiment. There are no significant associations between drought avoidance (A) or drought tolerance (B) with leaf number at the beginning of the experiment. Lower wilt VWC indicates greater drought avoidance while higher wilt survival indicating greater drought tolerance. Each point in both graphs is an individual. In the boxplots, box edges represent the interquartile range, the center line in the box is the median, and the whiskers represent 1.5 times less or greater than the interquartile range.

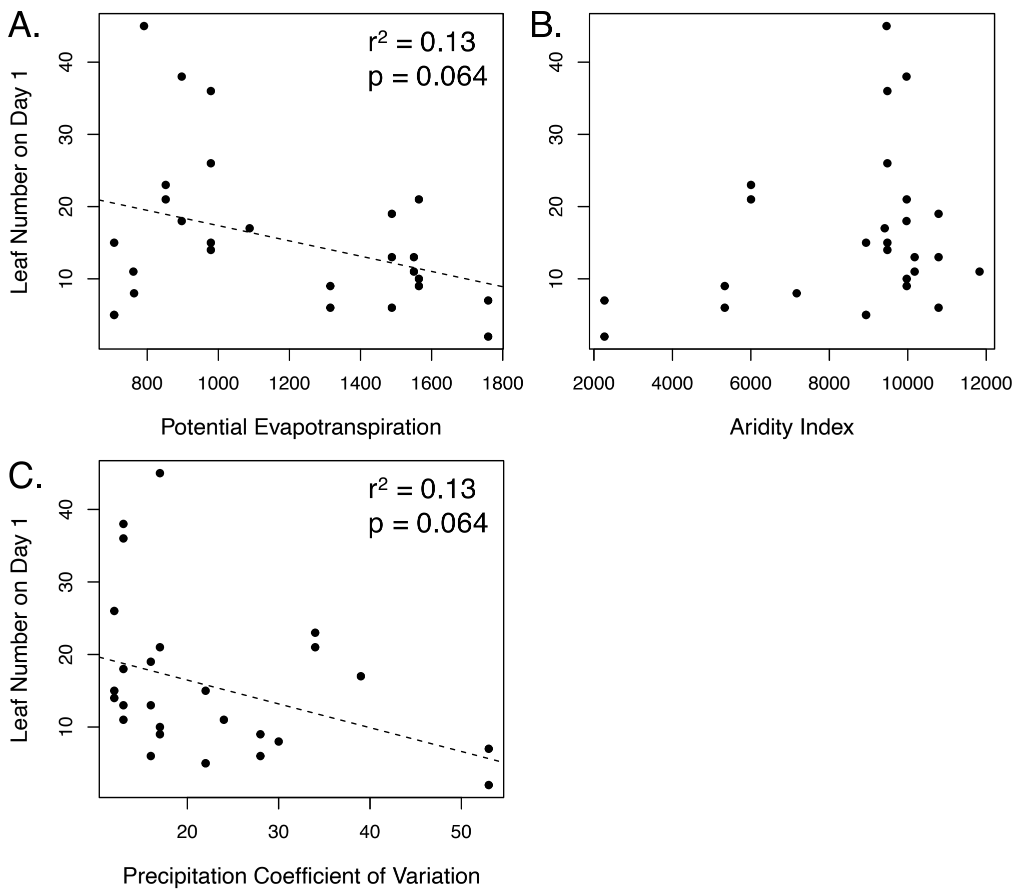


**Fig. S4** Associations between water-availability related variables of collection populations and leaf number at the start of the experiment.

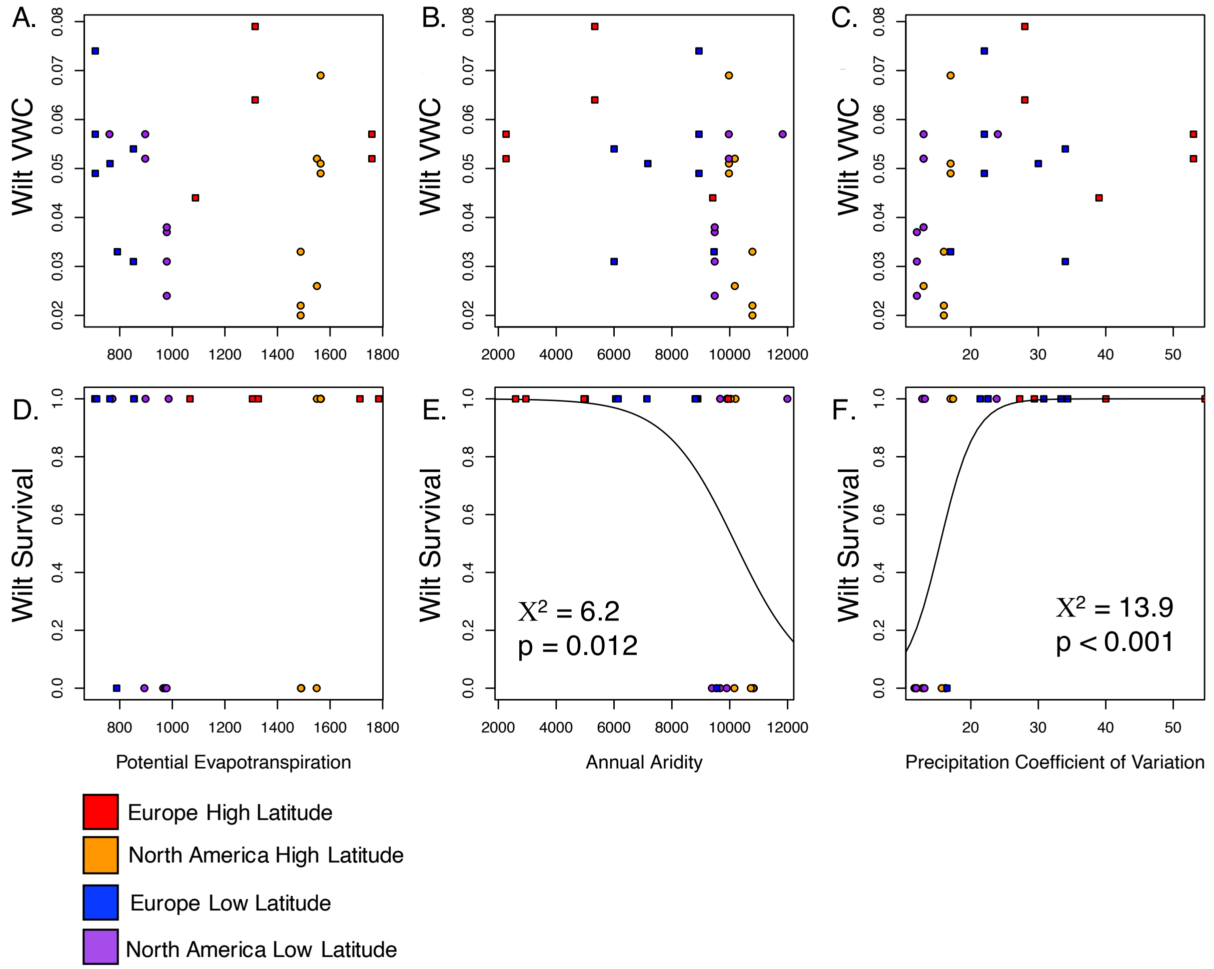


**Fig. S5** Associations between water-availability related variables of collection populations and drought avoidance and tolerance. Drought avoidance (A-C) or tolerance (D-F) relationships with Potential Evapotranspiration (A,D), Annual Aridity Index (B,E) or Annual Precipitation Coefficient of Variation (C,F). Solid lines indicate significant associations with associated test statistics and p-values on each plot. Note that these values are different than those in the main text or Table S3 as these models do not include leaf number at day 1 as a covariate – qualitative results remain the same. Colors and shapes represent differences among latitude and region contrasts.

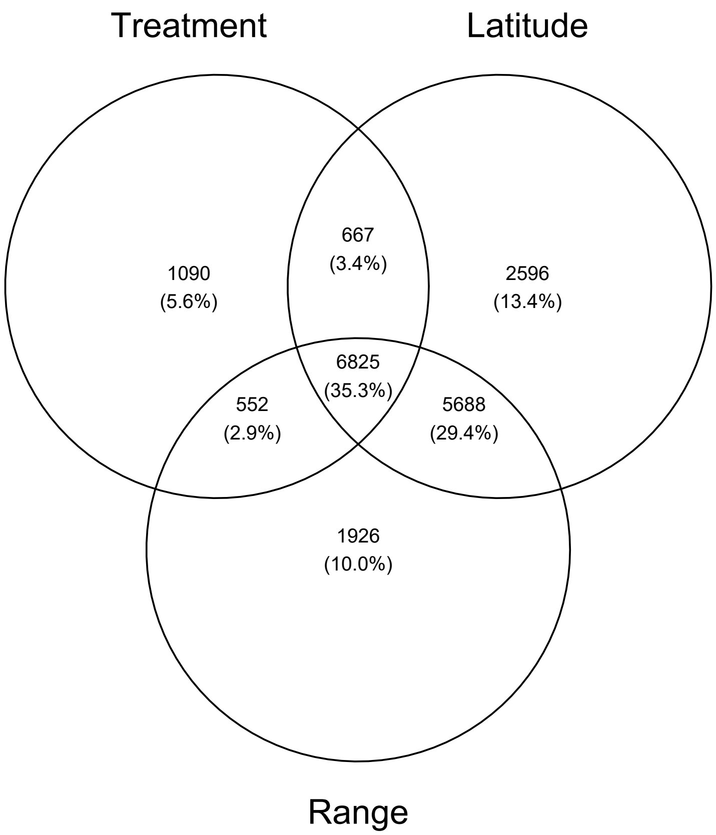


**Fig. S6** Venn diagram of differentially expressed genes across latitude, range and treatment contrasts. Numbers within overlapping circles represent genes that were differentially expressed in multiple contracts.

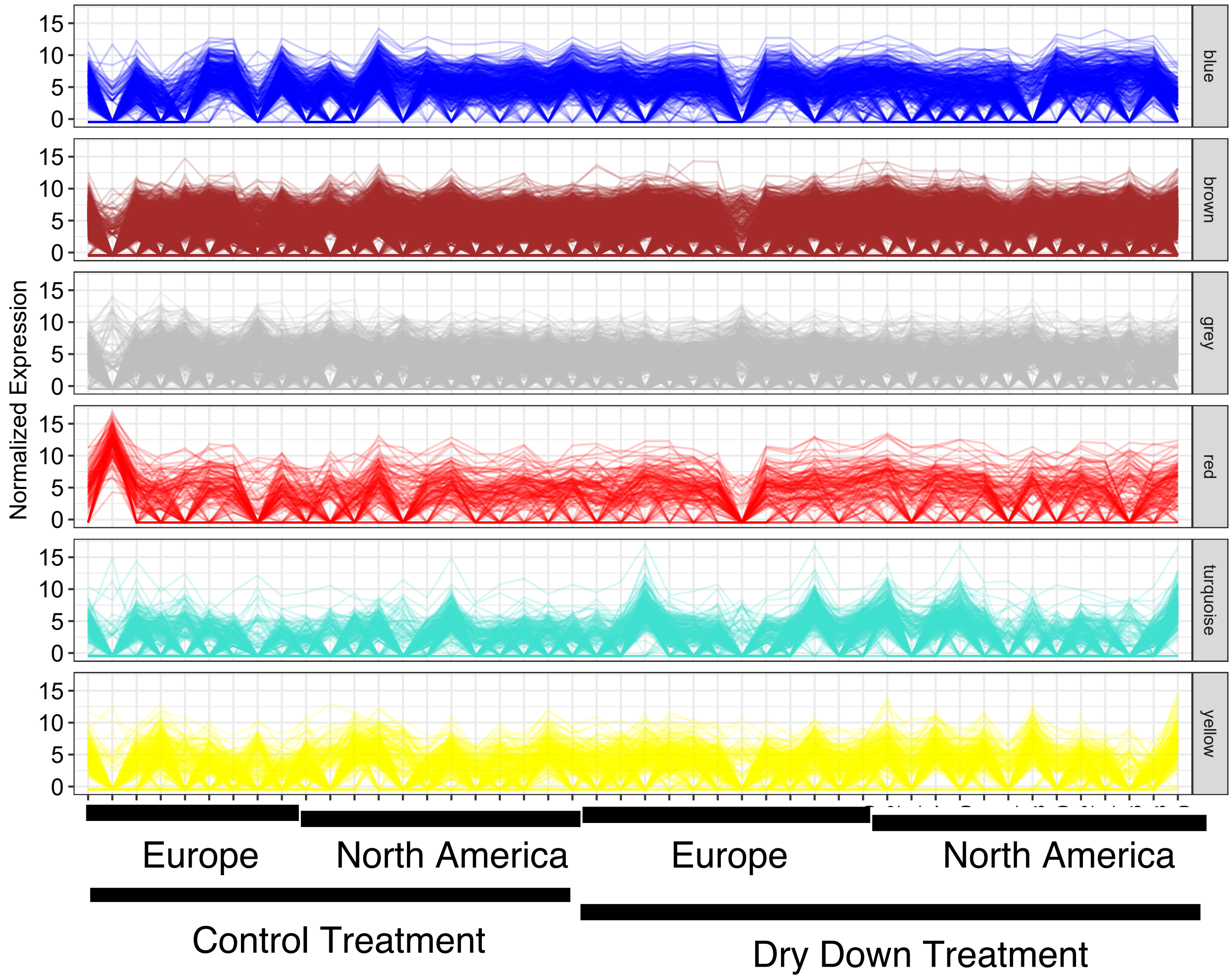


**Fig. S7**. Normalization expression profiles across samples for each module. Each tick on the x-axis is a different individual.

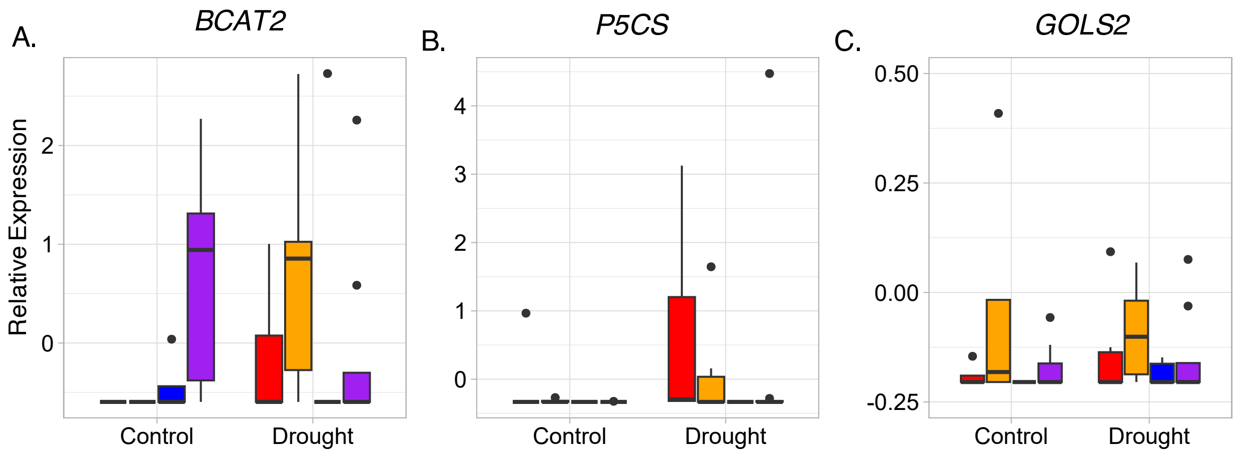


**Fig. S8** Relative expression across treatment, region, and latitude for candidate genes. Points and boxplots are colored by latitude and range: low latitude Europe = blue, high latitude Europe = red, low latitude North America = purple, High Latitude North American = orange. Each point represents a different individual.
